# *Isl1* Controls Axon Pathfinding of Retinal Ganglion Cells in the Binocular Visual Pathway

**DOI:** 10.1101/2025.03.03.640837

**Authors:** Shiona Biswas, Shuchun Li, Xiaoling Xie, Mei Xu, Cynthia Wen, Lin Gan

**Affiliations:** Department of Neuroscience & Regenerative Medicine, Medical College of Georgia, Augusta University, Augusta, GA 30912; James & Jean Culver Vision Discovery Institute, Medical College of Georgia, Augusta University, Georgia, GA 30912

**Keywords:** *Islet-1*, retina, optic chiasm, axon projection, bipolar vision, transcription factor

## Abstract

Axons of retinal ganglion cells (RGCs) face a choice to project ipsilaterally or contralaterally at the optic chiasm. This decision is crucial for the establishment of binocular vision in mammals. The transcription factor ZIC2 is well known as a key determinant of ipsilateral RGC identity, however the transcriptional programs controlling contralateral RGC identity have only begun to be elucidated more recently. Here we show that inactivation of the LIM-HD transcription factor ISL1 results in an expansion of the ZIC2^+^ ventrotemporal (VT) domain of the retina and an increase in ipsilateral projections from outside the VT retina. RNA-Seq, gene expression and CUT&Tag analyses show that ISL1 regulates RGC laterality through a set of unique downstream effectors as well as a few common ones, as other transcription factors. Our data also show that RGC axons misrouted at the midline in *Isl1*-null mutants could innervate the appropriate eye-specific regions of the dorsal lateral geniculate nucleus (dLGN) and superior colliculus (SC), in agreement with earlier studies that have shown that eye specific targeting of visual nuclei in the brain is independent of laterality decisions at the optic chiasm.

## Introduction

In species with binocular vision, different parts of each retina perceive the same visual field, which allows the animal to judge depth and distance of objects in relation to themselves and see the world in three dimensions (Nityananda and Read, 2017; Murcia-Belmonte and Erskine, 2019). Critical to the development of binocular vision is the establishment of the correct ratio of ipsilateral and contralateral retinal ganglion cells (RGCs) that segregate at the optic chiasm and project to visual targets on either the same side or the opposite side in the brain, respectively. In mice, which have laterally located eyes, only ∼3-5% of RGC axons project ipsilaterally (Rice et al., 1995; Jeffery and Erskine, 2005). Ipsilateral RGCs arise from the ventrotemporal (VT) retina from approximately embryonic day (E) 14 to 17.5, while contralateral RGCs arise from outside the VT retina from ∼E14 to E17, and from the VT retina from E17.5 to P0 (Herrera et al., 2003; Williams et al., 2006; Petros et al., 2008).

The zinc-finger transcription factor ZIC2 is expressed exclusively in the VT retina and acts as a key determinant of ipsilateral RGC fate by inducing expression of the receptor EPHB1 in ipsi-RGCs. The expression of EPHB1 in ipsi-RGCs results in their repulsion from the midline at the optic chiasm where the EPHB1 ligand EFNB2 (Ephrin-B2) is expressed by radial glial cells (Herrera et al., 2003; Williams et al., 2003; García-Frigola et al., 2008). Sonic hedgehog, secreted by contralateral RGCs at the optic chiasm, also repels ipsilateral RGCs from the midline (Peng et al., 2018).

Several receptor-ligand interactions are involved in the crossing of contralateral RGC axons at the midline. Vascular endothelial growth factor A (VEGFA), is expressed at the diencephalic midline where it acts as a chemoattractant for NRP1 (Neuropilin 1) expressed on contralateral RGCs, promoting their growth across the midline (Erskine et al., 2011). Interaction of NRCAM and PLXNA1 expressed on contralateral RGC axons with SEMA6D, PLXNA1 and NRCAM expressed on cells in the optic chiasm also promote contralateral RGC growth (Kuwajima et al., 2012). More recently, CXCL12 expressed in the meninges of the ventral diencephalon has been shown to drive axon growth toward the midline from CXCR4-expressing RGCs (Le et al., 2024).

ISL2, expressed in contralateral RGCs, regulates the development of only a specific subset of contralateral projections: those arising from the VT retina (Pak et al., 2004), while the SOXC family of transcription factors and POU3F1 have been shown to control development of the predominant set of contralateral RGCs: those arising from outside the VT retina (Kuwajima et al., 2017; Fries et al., 2023).

LIM-homeodomain (LIM-HD) proteins constitute a family of pleiotropic transcription factors that have roles in neuronal proliferation, fate specification, migration, axon pathfinding and dendritic arborization (Hobert and Westphal, 2000; Chou and Tole, 2019; Molina et al., 2023; Bose et al., 2024). Loss of function of the LIM-HD transcription factor ISL1 in flies and in spiral ganglia, sensory ganglia and cranial motor neurons of mice lead to defects in axon navigation (Thor and Thomas, 1997; Sun et al., 2008; Kim et al., 2016; Xu et al., 2024). In an earlier study, we have shown that ISL1 is expressed in developing RGCs immediately after they exit the cell cycle and is necessary for RGC axon growth, pathfinding and survival (Pan et al., 2008). In our current study, we show that upon deletion of ISL1, there is an expansion of the ZIC2^+^ RGC domain in the retina and an increased ipsilateral projection from regions of the retina that normally give rise only to contralateral RGCs. However, these mis-specified ipsilateral projections target regions of the dorsal lateral geniculate nucleus and superior colliculus that are normally innervated by contralateral RGC axons. Moreover, RNA-Seq and CUT&Tag analyses demonstrate that ISL1 regulates the expression of several transcription factors, cell adhesion molecules and axon guidance factors implicated in the development and axon pathfinding of contralateral versus ipsilateral projections.

## Methods

### Mice

Retina-specific inactivation of *Isl1* was achieved by breeding heterozygous *Isl1^lacZ/+^* or *Isl1^loxP/+^* mice with *Six3-Cre* mice and subsequently crossing with *Isl1^loxP/loxP^* mice. *Isl1^lacZ^* and *Isl1^loxP^*mouse lines were generated in our laboratory and have been described previously (Elshatory et al., 2007; Pan et al., 2008). *Six3-Cre* mice express Cre recombinase in the eye field and the ventral forebrain from E9 (Furuta et al., 2000) and have been used successfully as an effective retina-specific deleter (Mu et al., 2005; Pan et al., 2008). For timed matings, noon on the day a vaginal plug was observed was defined as embryonic day (E) 0.5. All analyses were conducted in mice of either sex. All animal procedures used in this study were approved by Institutional Animal Care and Use Committee (IACUC) at Augusta University.

### Immunohistochemistry and in situ hybridization

Heads of embryonic mice were fixed in 4% paraformaldehyde (PFA) in 0.1M phosphate-buffered saline (PBS), pH 7.2 at 4 °C for different times ranging from a few hours to overnight, after which they were transferred sequentially to 10%, 20% and finally 30% sucrose in PBS at 4 °C for cryopreservation and embedded in O.C.T. compound (Tissue-Tek, Sakura Finetek). Tissue was cryosectioned at a thickness of 14-20 µm, and immunostaining was performed as previously described (Dong et al., 2020). The following antibodies were used: anti-ß-galactosidase (lacZ) (Developmental Studies Hybridoma Bank 40-1a, 1:500), anti-ZIC2 (Sigma AB15392 discontinued, 1:200).

*In situ hybridization* (ISH) was performed as described previously (Biswas et al., 2023). Briefly, sections were fixed in 4% PFA, washed in 0.1M PBS, and treated with Proteinase K (1µg/ml). Hybridization was performed overnight at 70°C in hybridization buffer (4x SSC, 50% formamide, and 10% SDS) containing different antisense RNA probes. Post-hybridization washes were performed at 70°C in Solution X (2X SSC, 50% formamide, and 1% SDS). These were followed by washes in 2X SSC, 0.2X SSC, then TBS–1% Tween 20 (TBST). The sections were incubated in alkaline phosphatase-coupled anti-digoxigenin antibody (Roche) at 1:5,000 in TBST overnight at 4°C. The color reaction was performed using NBT/BCIP (Roche) in NTMT (100 mM NaCl, 100 mM Tris-pH 9.5, 50 mM MgCl2, and 1% Tween-20) according to the manufacturer’s instructions. *Isl1* and *Isl2* probes were described previously (Yang et al., 2003). Templates for *Cntn2*, *EphA5*, *EphrinA5*, *EphA7*, *EphB1*, *Nrcam* and *Zic2* were generated from plasmid DNA by restriction enzyme digestion. Probes for *Cxcr4*, *Gli1*, *Nrp1*, *Ptch1* and *Unc5c* were prepared by *in vitro* transcription using a kit from Invitrogen as per the manufacturer’s instructions. Templates for these probes were generated by PCR using specific primers, from E17.5 and E18.5 mouse retina cDNA. The sequence of the T7 polymerase promoter was added to the reverse primer sequences.

### Anterograde and Retrograde Labeling of RGCs

For visualization of the optic chiasm, small crystals of DiIC_18_ (Invitrogen) were placed over the optic disc of one eye in E18.5 embryos which had been fixed in 4% PFA in 0.1M PBS at 4°C for 1-2 days. After 14 days of incubation in PBS containing 0.1% sodium azide at 37°C, the diencephalon was dissected and imaged ventral side up.

For retrograde labeling of RGCs, rhodamine-conjugated dextran MW 6000 (Thermo Fisher Scientific) was placed in the anterior superior colliculus of P0 pups the heads of which had been fixed in 4% PFA at 4°C for 1-2 days. After incubation for 8 weeks in PBS containing 0.1% sodium azide at 37°C, retinae were dissected and flat-mounted in a Mowiol-based mounting medium for imaging.

Anterograde labeling of retinogeniculate projections using cholera toxin subunit B (CTB) were performed as described before (Murcia-Belmonte et al., 2019). Briefly, P13 pups were anesthetized, and the skin was cut open at the eyelid junction. Between 1-3 µl of CTB-Alexa Fluor 488 or CTB-Alexa Fluor 594 (Thermo Fisher Scientific) diluted in 1% DMSO were injected intravitreally. After 2 days, mice were anesthetized before transcardial perfusion with 4% PFA in PBS. Brains were fixed in 4% PFA overnight at 4°C and equilibrated in 30% sucrose before cryosectioning.

### RNA Sequencing (RNA-Seq)

Two repeats were performed for the strand-specific RNA-Seq experiments with retinae from mice at E13.5, and a pair of retinae were used for one sequencing sample. Retinae were flash frozen in Trizol and sent to Azenta Life Science (South Plainfield, NJ) for mRNA isolation and library preparation. The strand-specific RNA sequencing library was prepared using the Illumina NEBNext Ultra II Directional RNA Library Prep Kit according to the manufacturer’s instructions (New England Biolabs, Inc., Ipswich, MA), and PolyA selection for mRNA was used to remove rRNA. Sequencing was performed on the Illumina HiSeq 4000, and the depth was 20–30 million reads per sample. The Trimmomatic software (github.com/usadellab/Trimmomatic) was used to filter and trim minority low quality sequencing reads from the data set. The quality of sequence reads in FASTQ files was evaluated using the FastQC analysis. High quality sequence reads were aligned to the mouse mm10 reference transcriptome using HISAT2. The HISAT2 outputs were transformed with SAMtools and then fed into StringTie for transcript quantification. Subsequently, the count tables or matrices were input into the DESeq2 statistical package for determination of differential transcript expression between samples. The RNA-Seq data generated in this study have been deposited in the GEO database under accession number GSE290687.

Mutually exclusive and overlapping genes between datasets were identified using InteractiVenn (Heberle et al., 2015).

### CUT&Tag

The CUT&Tag experiment was performed using reagents and protocol provided by EpiCypher, Inc. with minor modifications as described previously (Xu et al., 2024). E13.5 retinae were isolated and nuclei were extracted immediately by incubating in NE Buffer for 10 min on ice. For each experiment, greater than 1 x 10^6^ nuclei were used. The nuclei were incubated with the activated concanavalin A-coated magnetic beads (11 μl/sample, EpiCypher #21-1401) at room temperature for 10 min. The nuclei-beads were collected using magnetic stand and resuspended in 50 μl ice cold Antibody Buffer. 0.5 μg antibody against ISL1 (rabbit, AB4326, Millipore; rabbit, ab20670, Abcam) was added to each sample and mixed on a rotator at 4°C overnight or at room temperature for 2 h. The antibody-bound nuclei-beads were isolated and resuspended in 50 μl of ice cold Digitonin 150 Buffer. 0.5 μg of anti-rabbit IgG secondary antibody (EpiCypher #13-0047 and #13-0048) was added and samples were incubated for 30 min at room temperature on a rotator. After two washes in ice cold Digitonin 150 Buffer, nuclei-beads were resuspended in 50 μl of ice cold Digitonin 300 Buffer, 2.5 μl of CUTANA pAG-Tn5 solution (EpiCypher #15-1017) was added, and nuclei-beads were incubated for 1 h at room temperature on a rotator. The samples were again washed twice in ice cold Digitonin 300 Buffer, resuspended in 125 μl of cold Tagmentation Buffer, and incubated at 37°C for 1 h. Then, 2.5 μl of 0.5 M EDTA, 1.25 μl of 10% SDS and 1.1 μl of 20 mg/ml Proteinase K were added to each sample and samples were incubated at 70°C for 10 min or 55°C for 30 min. After this, DNA fragments were purified and collected in 30 μl of ddH2O using Cycle Pure Kit (Omega, D6492-02) following the manufacturer’s instruction. To prepare the library, 21 μl of DNA fragments were transferred to a PCR tube, to which 2 μl of barcoded i5 and i7 primers and 25 μl CUTANA High Fidelity 2x PCR Master Mix (EpiCypher, #15-1018) were added. PCR was performed using the following conditions: 72°C for 5 min, 98°C for 30 s, and 20 cycles of 10 s at 98°C and 10 s at 63°C, followed by an extra 1 min extension at 72°C. The PCR product was analyzed on an Agilent Fragment Analyzer. Combined DNA libraries were cleaned up using AMPure XP beads (#A63880, Beckman Coulter) following the manufacturer’s instructions. Each pooled DNA library of 10 samples was resuspended into 40–80 μl water and quantified using the NanoDrop (Thermo Scientific). Library fragments were sent for sequencing commercially at Azenta Life Science. Sequencing was performed on Hiseq 4000 (Illumina) with 2 x 150 paired-end reads and a depth of 10 million reads per sample. The CUT&Tag with anti-ISL1 antibody were performed in three replicates. Raw data of each replicate was analyzed following the CUT&Tag Data Processing and Analysis Tutorial (https://yezhengstat.github.io/CUTTag_tutorial/). Briefly, raw data was pre-processed for quality control using FastQC. Qualified peaks were aligned to the mouse mm10 reference transcriptome using Bowtie2. Peaks were called using MACS2 and SEACR. Peak annotation and functional analysis were performed using ChIP seeker and visualized by IGV. The accession number for the CUT&Tag-Seq data is GSE290824

### Experimental Design and Statistical Analysis

For Figure 1, the fluorescence intensity within a designated area of the same size in the ipsilateral and contralateral optic tracts was calculated using ImageJ (NIH). Fluorescence was computed as the corrected pixel intensity difference between the “integrated density” of the selected area and the mean fluorescence of background readings from areas of the same size. The ipsilateral index was defined as the ratio of the fluorescence intensity in the ipsilateral optic tract to the sum of the fluorescent intensity in the ipsilateral optic tract and the contralateral optic tract. Statistical analysis was performed using GraphPad Prism software version 10. In Figure 4B, three differentially expressed genes could not be represented in the volcano plot as their log2-fold change (Eef1akmt4-ece2: 21.64406851, Havcr1: −23.35119895) or P value (Gli1: 8.05061310590541E-202) were too high.

**Figure 1.**
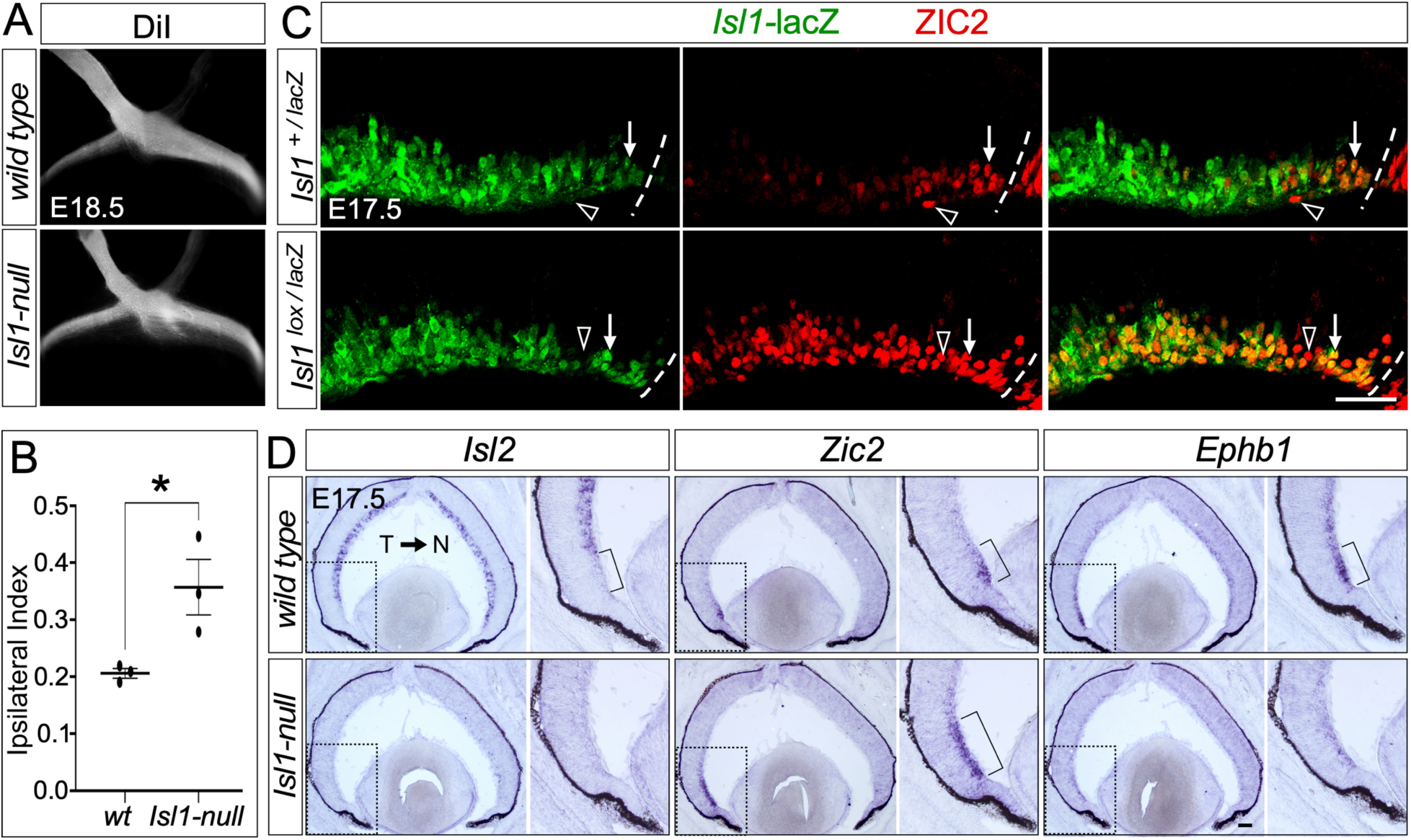
Loss of Isl1 from the retina results in misrouting of ipsilateral RGC projections and expansion of the retinal ipsilateral domain. ***A,*** Ventral views of the optic chiasm labeled from one eye at E18.5 reveal a thicker ipsilateral tract relative to contralateral in *Isl1-null* mutants compared to controls. ***B,*** Quantification of fluorescence intensity in ipsilateral optic tract, divided by the sum of the fluorescence intensity in the ipsilateral optic tract and the contralateral optic tract, shows an increased ipsilateral index in *Isl1-null* mutants. Unpaired two-tailed Student’s t test, p = 0.0374; n = 3 for each genotype. ***C,*** lacZ and ZIC2 immunostaining at E17.5 show that many ZIC2^+^ cells co-express ISL1 (arrows) in control retina, while in *Isl1-null* retina, ZIC2 can be found in more centrally located RGCs outside the ventrotemporal retina, including cells from which *Isl1* was deleted (arrows). Arrowheads in control and mutant retinae indicate RGCs that express only ZIC2. ***D,*** Control and *Isl1-null* sections at E17.5 displaying expression of *Isl2, Zic2 and EphB1*. In *Isl1-nulls*, *Isl2* is dramatically reduced, *EphB1* expression much weaker, while the *Zic2* expression domain is expanded centrally relative to WT. Brackets in each inset indicate the ventrotemporal retina. CMZ, ciliary margin zone; L, lens; NR, neural retina; N, nasal region; T, temporal region. Scale bars: 100 µm.

## Results

### Loss of ISL1 results in an expansion of the ZIC2^+^ ventrotemporal domain and an increase in ipsilateral projections

We have previously demonstrated that retina-specific *Isl1* knockout mice display axon growth defects at the optic chiasm (OC), and the optic nerve and tracts are thinner due to a reduction in the number of RGCs (Pan et al., 2008). To investigate further if there was misrouting of ipsilateral or contralateral projections at the OC, we used the lipophilic dye DiI to anterogradely label RGC axons from one eye of E18.5 embryos. In contrast to wild-type embryos with a prominent contralateral projection, *Isl1-null* mice displayed a significant increase in the proportion of axons projecting ipsilaterally (Fig. 1*A*). Quantification of the fluorescence intensity in similarly sized regions of the ipsilateral and contralateral optic tract revealed an increased ipsilateral index in *Isl1-null* mice (Fig. 1*B*). We next determined whether *Isl1* is expressed selectively in contralaterally projecting RGCs. Immunolabeling of control *Isl1-lacZ* (*Isl1^loxP/lacZ^*) and *Isl1-null* (*Isl1^loxP/lacZ^*; *Six3-Cre*) retinae at E17.5 with anti-lacZ and anti-ZIC2 antibodies revealed that in controls, *Isl1-*lacZ is expressed throughout the ganglion cell layer, but at a lower level in the ventrotemporal (VT) retina, where ZIC2^+^ RGCs reside: *Isl1-*lacZ is present in many (arrows) but not all (arrowhead) ZIC2^+^ RGCs (Fig. 1*C*). In contrast, there is an expansion of ZIC2^+^ RGCs into more central regions of the retina in *Isl1-*nulls, including in RGCs from which *Isl1* was deleted (arrows in Fig. 1*C*). Examination of mRNA expression of the contralateral RGC marker *Isl2* (Pak et al., 2004), ipsilateral RGC marker and fate determinant *Zic2* (Herrera et al., 2003; García-Frigola et al., 2008), and the axon guidance receptor EPHB1 that mediates the repulsive response of EPHB1-expressing ipsilateral RGC axons with its ligand ephrin B2 (EFNB2) expressed at the OC midline (Williams et al., 2003), showed that in *Isl1-*nulls, the expression of *Isl2* was abolished, the *Zic2* expressing domain expanded centrally, consistent with our immunostaining data, and interestingly, *EphB1* expression was decreased (Fig. 1*D*). Taken together, our data indicate that *Isl1* is expressed in both contralateral and ipsilateral RGCs, and ISL1 is required for the expression of *Isl2* and *EphB1* but represses ZIC2 expression.

### *Isl1* inactivation leads to an increase in ipsilateral projections from RGCs in the contralateral retina

To determine where in the retina the increased ipsilaterally projecting RGCs in *Isl1*-null mutants originate from, we performed unilateral retrograde rhodamine dextran labeling from the anterior superior colliculus (SC) at P0 and examined retinal flat mounts of the eye ipsilateral to dextran injected SC (Fig. 2*A*). Unlike in controls (Fig. 2*B*,*C*, top), where labeled RGCs were ipsilateral RGCs and found exclusively in the ventrotemporal retina, labeled RGCs were not limited to the VT area but were throughout the retina in the *Isl1*-null mutants (Fig. 2*B,C*, bottom). Additionally, the density of RGCs appeared lower in the VT area of *Isl1-*nulls, which could be explained by the fact that *EphB1* expression is reduced in these mice (Fig. 1*D*).

**Figure 2.**
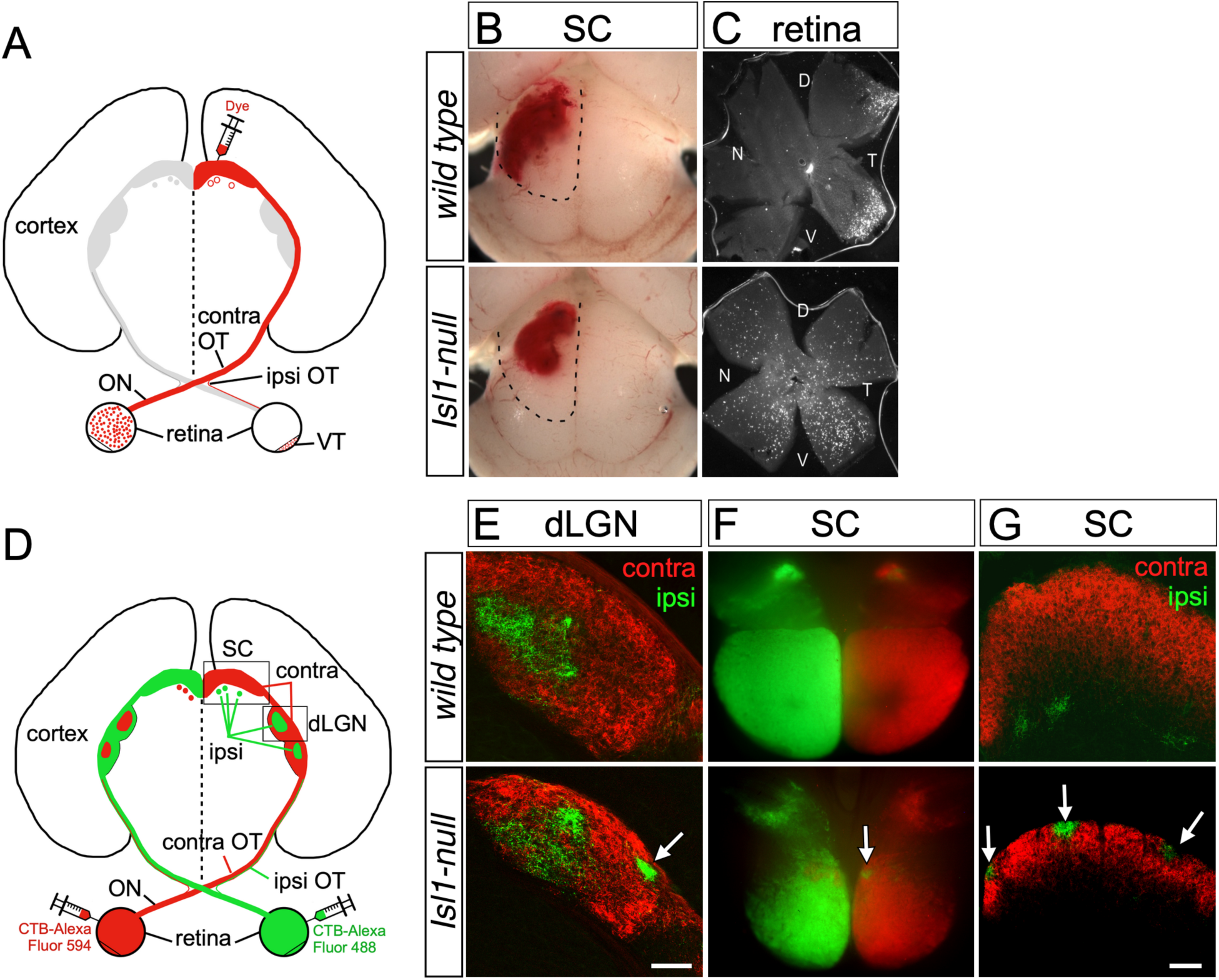
Eye specific targeting of brain nuclei is independent of laterality decisions at the midline upon loss of *Isl1*. ***A,*** Schematic drawing of unilateral retrograde tracing of RGCs. ***B,*** Rhodamine dextran was injected in the superior colliculus at P0, as indicated. ***C,*** Retinae ipsilateral to the rhodamine dextran-injected SC were analyzed. Ipsilaterally projecting RGCs are restricted to the VT retina in controls but can be found in all four quadrants of the *Isl1-null* retina. ***D,*** Schematic drawing of anterograde labeling of retinogeniculate and retinocollicular projections. At P13, CTB-Alexa Fluor 488 was injected into one eye and CTB-Alexa Fluor 594 into the other and retinal axons were visualized in the dLGN and SC of controls and *Isl1-nulls* at P15. ***E,*** In control mice (upper panel), coronal sections of the dLGN show ipsilateral projections (green) forming a patch in the dorsomedial dLGN surrounded by contralateral projections (red). In *Isl1-null* mice (lower panel), ipsilateral axons can be seen within a region that is normally innervated by contralateral axons (arrow), in addition to some ipsilateral projections to the dorsomedial dLGN. ***F*** and ***G,*** In controls (upper panels), ipsilateral axons can be found in small patches in the deeper SO, while contralateral axons innervate the more superficial SGS as seen in whole mounts of the SC (***F***), and coronal sections (***G***). In *Isl1-nulls* (lower panels), ipsilateral axons can be found in the SGS in SC whole mounts (***F***), and coronal sections (***G***) (arrows). D, dorsal region; V, ventral region, N, nasal region; T, temporal region, dLGN, dorsal lateral geniculate nucleus; SC, superior colliculus; SGS, stratum griseum superficiale; SO, stratum opticum; CTB, cholera toxin subunit B. Scale bars: 100 µm.

Two of the most important brain nuclei that RGCs project to are the dorsal lateral geniculate nucleus (dLGN), which conveys information to the visual cortex for image forming, and the SC, which processes visual input for non-image forming functions, such as coordination of head and eye movements and fear and avoidance behaviors (Assali et al., 2014; Murcia-Belmonte and Erskine, 2019). To investigate the effect of *Isl1* inactivation on RGC projections to LGN and SC, we injected cholera toxin subunit B (CTB)-Alexa Fluor 488 into one eye and CTB-Alexa Fluor 594 into the other eye of mice at postnatal day 13 (P13) and examined the dLGN and SC at P15, a time point at which eye specific segregation of retinal projections is largely complete (Godement et al., 1984; Jaubert-Miazza et al., 2005). In the dLGN of controls, ipsilateral axons were present in a core in the dorsomedial region, surrounded by contralateral axons (Fig. 2*E*, top). In the dLGN of *Isl1-*null mutants however, while ipsilateral axons were seen in the dorsomedial region, they also projected to the normally contralateral axon-recipient region of the dLGN (arrow, Fig. 2*E*, bottom). In the SC of controls, contralateral axons were localized in the most superficial layer of the SC, the stratum griseum superficiale (SGS), while ipsilateral axons were found in an underlying layer, the stratum opticum (SO) (Fig. 2*F,G*, top) (Godement et al., 1984; Assali et al., 2014). In contrast, in *Isl1-*null mutants, ipsilateral axons could be seen amidst contralateral axons in the SGS (arrows, Fig. 2*F,G*, bottom). Thus, RGC axons that originated from contralateral domain but were misrouted at the midline could still innervate the appropriate eye-specific regions of the dLGN and the SC. These results indicate that broad target recognition is intrinsically encoded in RGCs and is related to their retinal position, which is in agreement with earlier studies (Rebsam et al., 2009; García-Frigola and Herrera, 2010).

### The expression of Eph-ephrin axon guidance factors is unaltered in *Isl1*-null mutants

Members of the Eph family of receptor tyrosine kinases and their ephrin ligands play important roles in the development of the retinogeniculate and retinocollicular projection. EphA receptors and their ligands are expressed in opposing nasotemporal gradients in the retina, opposing ventrolateral and dorsomedial gradients in the dLGN and vLGN, and opposing anteroposterior gradients in the SC (Cang and Feldheim, 2013). Repulsive EphA/Ephrin-A interactions result in an ordered topographic patterning of retinal projections in these target structures in the brain (Feldheim et al., 1998, 2000; Cang and Feldheim, 2013). Therefore, we examined the expression of these signaling factors. In the RGC layer of the retina, *EphA5* is expressed in a temporal (high)-to-nasal (low) gradient, while *EphrinA5* is expressed in a nasal (high)-to-temporal (low) gradient (Fig. 3*A*,*B*, upper panels) (Feldheim et al., 2000; Cang and Feldheim, 2013), a pattern that was unchanged in the *Isl1-*null mutants (Fig. 3*A*,*B*, lower panels). The ventrolateral (high)-to-dorsomedial (low) gradient of expression of *EphrinA5* in the dLGN and vLGN and the counter-gradient of *EphA7* (Fig. 3*C*,*D*, upper panels) (Feldheim et al., 1998; Pfeiffenberger et al., 2005) was also unchanged in *Isl1-*null mutants (Fig. 3*C*,*D*, lower panels), as was the anterior (high)-to-posterior (low) gradient of *EphA5* and *EphA7* expression in the SC (Fig. 3*E*,*F*) and the opposing gradient of *EphrinA5* expression (Fig. 3*G*) (Feldheim et al., 2004; Pfeiffenberger et al., 2005). These results suggest that defective bifurcation and projection of RGC axons in *Isl1*-null retinae were not caused by changes in the Eph-ephrin signaling pathway.

**Figure 3.**
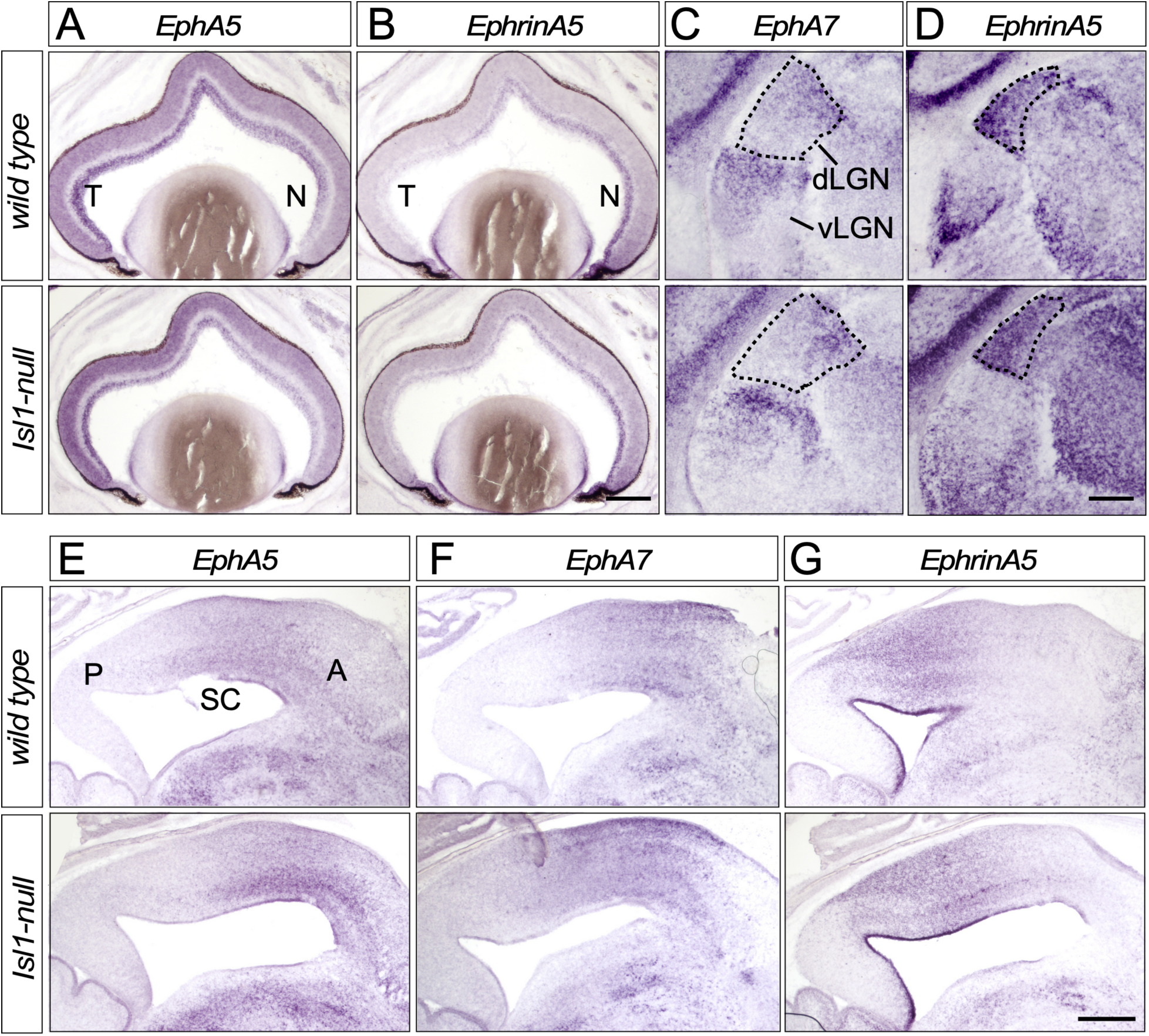
Expression of Eph/Ephrin signaling factors are unchanged upon loss of *Isl1*. ***A*** and ***B,*** At P0, *EphA5* (***A***) and *EphrinA5* (***B***) expression patterns are unchanged in the *Isl1-*null retina (lower panels), compared to control (upper panels). ***C*** and ***D,*** Coronal sections of the dLGN and vLGN at P0 reveal unchanged expression of *EphA5* (***C***) and *EphrinA5* (***D***) in *Isl1-*null (lower panels), compared to control (upper panels). ***E***-***G,*** Sagittal sections of the SC at P0 reveal unchanged expression of *EphA5* (***E***), *EphA7* (***F***) and *EphrinA5* (***G***), in the *Isl1*-null (lower panels) compared to WT (upper panels). N, nasal region; T, temporal region; A, anterior; P, posterior; dLGN, dorsal lateral geniculate nucleus; vLGN, ventral lateral geniculate nucleus; SC, superior colliculus. Scale bars: ***B,*** 250 µm; ***D,*** 100 µm; ***G,*** 500 µm.

### *Isl1* functions cell-autonomously in RGCs to regulate the intrinsic projection properties of RGC axons

*Isl1* is also expressed in parts of the ventral forebrain and we have previously used the *Six3-Cre* mouse line to achieve deletion of *Isl1* in this region (Furuta et al., 2000; Elshatory et al., 2007). We next examined the expression of crucial axon guidance factors in both the retina and diencephalic midline of *Isl1-null* mutants. Contactin-2/TAG1 is a cell adhesion molecule required for the fasciculation of RGC axons during development (Chatzopoulou et al., 2008). The expression of *Cntn2* is also significantly reduced in the *Isl1-null* mutant retina (Fig. 4*A*). The expression of the cell adhesion molecule NRCAM is required in both contralateral RGCs and radial glia at the optic chiasm for contralateral RGCs to cross (Williams et al., 2006; Kuwajima et al., 2012). We found that *Nrcam* expression is markedly reduced in the retina (Fig. 4*B*) but unchanged at the diencephalic midline (Fig. 4*C*) of *Isl1-*null mice. The ligand EFNB2 is expressed on radial glia at the optic chiasm and binds to its receptor EPHB1 which is expressed by ipsilateral RGCs, resulting in their repulsion away from the midline and into the ipsilateral optic tract (Williams et al., 2003). *Efnb2* expression is unchanged at the midline of *Isl1*-null mutants (Fig. 4*D*). In addition, we could not detect ISL1 protein at the optic chiasm of control or *Isl1-*null mice (Fig. 4*E*). Taken together, these data show that misrouting of RGC axons in *Isl1-*null mutants was largely caused by changes intrinsic to RGCs and not due to non-cell autonomous mechanisms at the OC.

**Figure 4.**
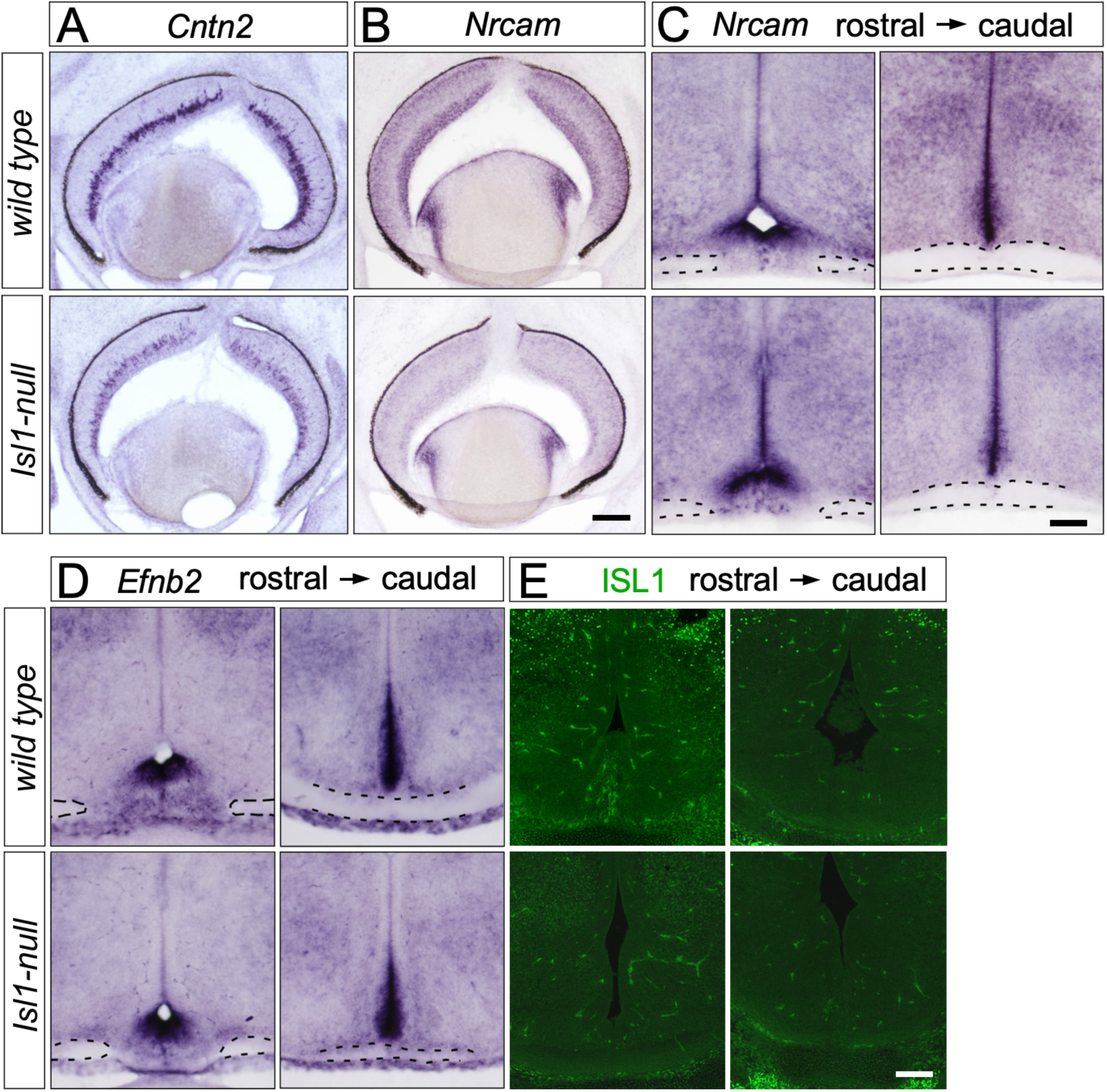
Expression of RGC axon guidance factors are changed in the retina but not at the diencephalic midline of *Isl1-null* mutants. ***A,*** *Cntn2* expression is decreased in the *Isl1-null* retina at E14.5. ***B,*** *Nrcam* expression is downregulated in the *Isl1-null* retina at E14.5. ***C,*** *Nrcam* expression is unchanged at the diencephalic midline of *Isl1-null* mutants compared to controls, as seen in two different rostro-caudal levels. ***D,*** *Efnb2* expression is unchanged at the diencephalic midline of *Isl1-null* mutants compared to controls, as seen in two different rostro-caudal levels. ***E,*** Anti-ISL1 immunostaining at E17.5 shows no detectable ISL1 expression at the optic chiasm. Dashed lines indicate retinal axons. Scale bars: 100 µm.

### Loss of *Isl1* results in the down-regulation of genes enriched in contralateral RGCs

To get an overall picture of the transcriptomic changes upon loss of *Isl1*, we performed RNA-Seq on control and *Isl1-*null retinae at E13.5 (Fig. 5*A*). We found 576 downregulated genes and 175 upregulated genes in the *Isl1-*null relative to control, using a P_adj_ < 0.05. Among the genes that displayed downregulation (Fig. 5*B*) were *Isl2* which regulates the late-born contralateral RGC projection from the VT retina (Pak et al., 2004), and which is consistent with our ISH data at E17.5. Other downregulated genes include axon growth receptors and guidance cues such as *Cxcr4*, which is required for contralateral axon outgrowth at the optic chiasm (Le et al., 2024), *Nrp1* (Neuropilin1), a receptor that is expressed in and required for contralateral RGCs to cross at the optic chiasm (Erskine et al., 2011; Kuwajima et al., 2012), *Shh* (Sonic hedgehog), which is secreted by contralateral RGCs at the optic chiasm to repel ipsilateral RGCs (Peng et al., 2018), and *Cntn2.* The downregulation of *Cntn2* is also consistent with our *in situ hybridization* data. The POU-homeobox gene *Pou4f1* (aka *Brn3a*) was also significantly downregulated; transcripts for *Pou4f1* were highly enriched in contralateral RGCs in two studies that separately analyzed the transcriptomic signatures of ipsilateral and contralateral RGCs (Wang et al., 2016; Fernández-Nogales et al., 2022). (Fernández-Nogales et al., 2022) additionally demonstrated that POU4F1 is essential for midline crossing as disruption of its function at E13.5 led to a reduced number of axons crossing at the optic chiasm. Relevant to these findings, our earlier study has shown a marked reduction in the expression of *Shh* and *Pou4f1* mRNA in the *Isl1-null* retina at E14.5 and that ISL1 binds to the promoters of *Shh*, *Pou4f1* and *Isl2* via chromatin immunoprecipitation (ChIP) in retinae (Pan et al., 2008).

**Figure 5.**
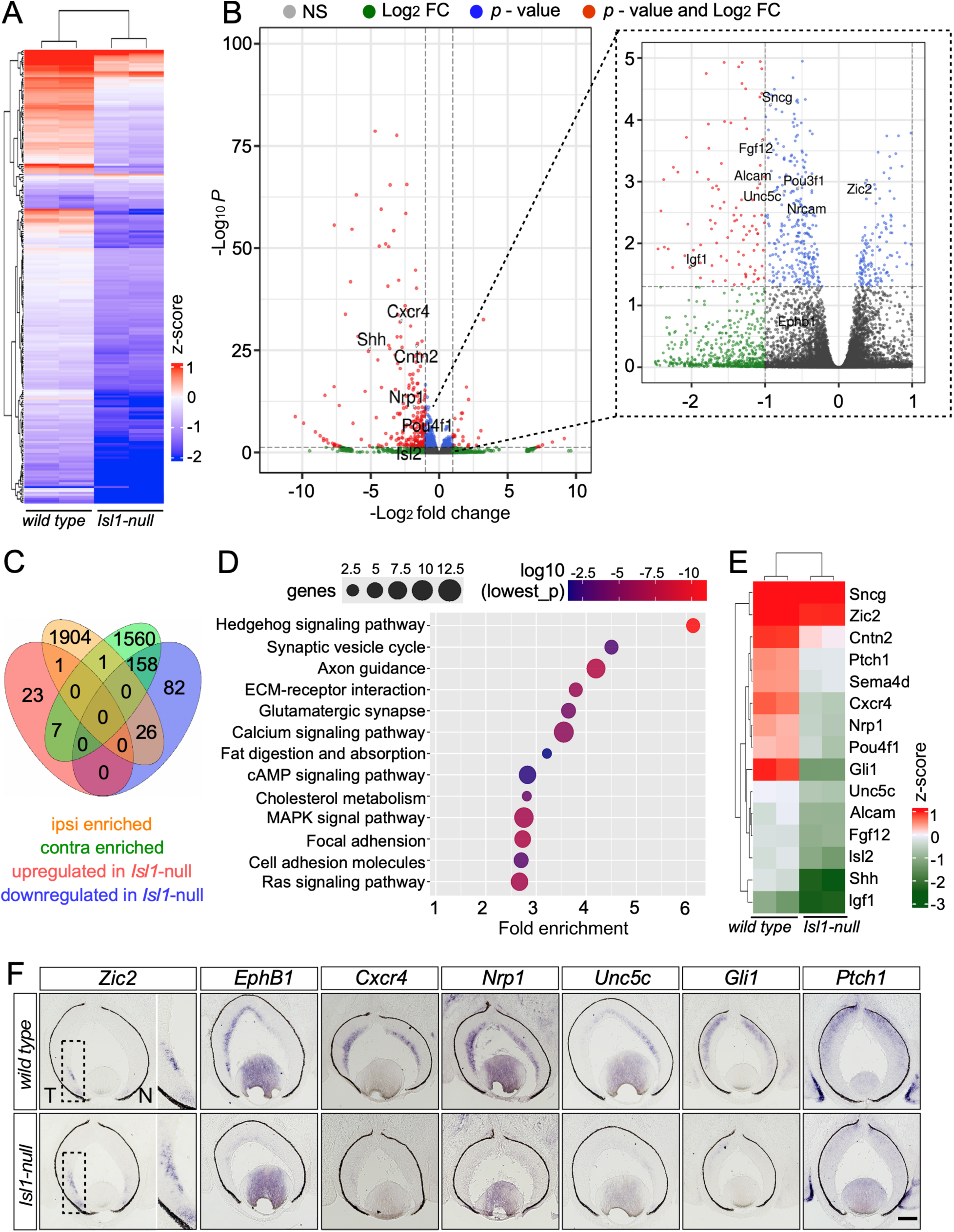
Transcriptomic analysis of the *Isl1-null* retina. ***A,*** Overall picture of differentially expressed genes revealed by RNA-Seq in the *Isl1-null* retina compared to control at E13.5. ***B,*** Volcano plot illustrating downregulated and upregulated genes in the *Isl1-null*, with genes of interest highlighted. Inset on the right is a higher magnification view of the boxed region on the left. ***C,*** Venn Diagram showing the number of DEGs found in *Isl1-null* RNA-Seq that overlap with genes enriched in ipsilateral or contralateral RGCs at E16.5 as shown in an already published RNA-Seq dataset (Fernández-Nogales et al., 2022). Ipsi enriched: genes significantly enriched in ipsilateral RGCs, contra enriched: genes significantly enriched in contralateral RGCs. ***D,*** KEGG analysis of differentially expressed genes in the *Isl1-null* retina. ***E,*** Heat map of selected genes that are differentially expressed genes in the *Isl1-null* retina. ***F,*** ISH validation of RNASeq results for selected genes that are differentially expressed in the *Isl1-null* retina at E14.5. N, nasal region; T, temporal region. Scale bar: 100 µm.

Other significantly downregulated genes were the cell adhesion molecule *Alcam*, important for RGC axogenesis (Thelen et al., 2012), *Igf1* and *Fgf12* which are enriched in the transcriptome of contralateral RGCs and are expressed outside the Zic2^+^ VT retina (Wang et al., 2016; Fernández-Nogales et al., 2022). Among the genes that were significantly downregulated but with a fold change < 1, were *Nrcam*, *Unc5c*, *Sncg* and *Pou3f1* (Fig. 5*B*, right). *Nrcam* expression is also reduced in the *Isl1-null* retina (Fig. 4*A*). *Unc5c* (UNC5C) is an axon guidance receptor required for the growth of retino-retinal axons and its expression is repressed by ZIC2 in ipsilateral RGCs (Murcia-Belmonte et al., 2019; Fernández-Nogales et al., 2022). *Sncg* (SNCG, *γ*-synuclein) is a transcriptional target of POU4F1 and required for midline crossing of contralateral RGCs (Fernández-Nogales et al., 2022). A recent study reported that the transcription factor *Pou3f1* (POU3F1) is expressed in contralateral RGCs and required for their midline crossing (Fries et al., 2023). *Zic2* is also upregulated in the *Isl1-*null, consistent with our immunostaining and ISH data at E17.5 (Fig. 1*C,D*). Additionally, our RNA-Seq data also shows a 0.47-fold downregulation of *Ephb1* in the *Isl1-*null although this change is not significant.

We next cross-referenced our RNA-Seq dataset with a previously published RNA-Seq dataset that examined differentially expressed genes between ipsilateral and contralateral RGCs at E16.5 (Fernández-Nogales et al., 2022) using a cutoff of log_2_fold change > 1, and P_adj_ < 0.05 for both. We found that 158 of the 266 downregulated genes in the *Isl1-*null retina, i.e., ∼60% were enriched in contralateral RGCs (Fig. 5*C*). These included *Isl2*, *Nrp1*, *Shh*, *Cntn2*, *Pou4f1*, *Igf1* and *Fgf12* discussed above, and additional genes such as *Ngfr*, *Pou6f2* and *Rgs4.* Only a small proportion (∼10%) of downregulated genes in the *Isl1-null* dataset were enriched in ipsilateral RGCs. These included the Shh signaling pathway effectors *Gli1* and *Ptch1* and the axon guidance receptor *Sema4d*. Interestingly, this list also included *Slc6a4* which codes for the serotonin transporter (SERT). *Slc6a4,* a transcriptional target of ZIC2, is not required for RGC axons to project correctly at the midline but is essential for the refinement of ipsilateral RGC projections in the superior colliculus (García-Frigola and Herrera, 2010). We further analyzed the DEGs in the *Isl1-null* retina using KEGG Pathway analysis (Fig. 5*D*). This revealed multiple dysregulated genes involved in axon guidance (*Cxcr4*, *Nrp1*, *Shh*, *Ptch1* and *Sema4d*), the Hedgehog signaling pathway (*Shh*, *Ptch1* and *Gli1*), and cell adhesion molecules (*Alcam* and *Cntn2*) as shown in the heat map of selected differentially expressed genes (Fig. 5*E*). Using *in situ hybridization,* we have already shown a change in expression of some of these, such as *Cntn2* at E14.5 and *Zic2* and *Isl2* at E17.5 (this study), and *Pou4f1*, *Isl2* and *Shh* at E14.5 (Pan et al., 2008). We used ISH to validate the RNA-Seq results for some more of these genes: *Cxcr4*, *Nrp1*, *Unc5c*, *Gli1* and *Ptch1,* and confirmed the change in expression of *Zic2* and *EphB1* at E14.5 (Fig. 5*F*). In summary, our RNA-Seq data suggest that ISL1 directly or indirectly regulates the expression of several cell adhesion molecules, axon guidance receptors and transcription factors required for contralateral RGCs to cross the diencephalic midline.

### ISL1 directly regulates genes essential for the identity and axon guidance of ipsilateral and contralateral RGCs

To determine how ISL1 regulates the DEGs identified by RNA-Seq analysis, we identified genomic regions occupied by ISL1 in the retina using CUT&Tag approach (Kaya-Okur et al., 2019) and data from duplicate experiments were combined (Fig. 6*A*). The locations of ISL1-bound peaks showed a significant enrichment at the proximity of transcriptional start sites (TSS) (Fig. 6*B,C*), suggesting a role in initiating transcription. Genomic feature annotation showed that almost half of ISL1 (∼40%) peaks were enriched within 1 kb of promoter regions, which is consistent with its known role, while ∼20% were in distal intergenic regions (Fig. 6*D*). To identify potential genes directly regulated by ISL1, we first filtered the peak regions for predicted ISL1-binding sites by Bedtools intersect and identified a total of 2,779 ISL1-bound genes (Fig. 6*E*). By integrating these genes with the DEGs identified by RNA-Seq analysis, we obtained 137 downregulated and 40 upregulated genes bound by ISL1 (Fig. 6*E*). The CUT&Tag peaks of ISL1 were further examined by the representative Integrative Genomics Viewer (IGV) genome track snapshots to reveal the binding sites of ISL1 at the DEGs identified by RNA-Seq analysis. We found ISL1 binding at cis-regulatory elements of genes encoding cell adhesion molecules, axon guidance receptors and transcription factors required for the development of ipsilateral and contralateral RGCs, including *Cxcr4*, *Cntn2*, *Alcam*, *Nrcam*, *Zic2* and *Isl2 (*Fig. 5*F*). Thus, these discoveries indicate that ISL1 directly binds to and regulate the expression of genes essential for the identity and axon guidance of ipsilateral and contralateral RGCs.

**Figure 6.**
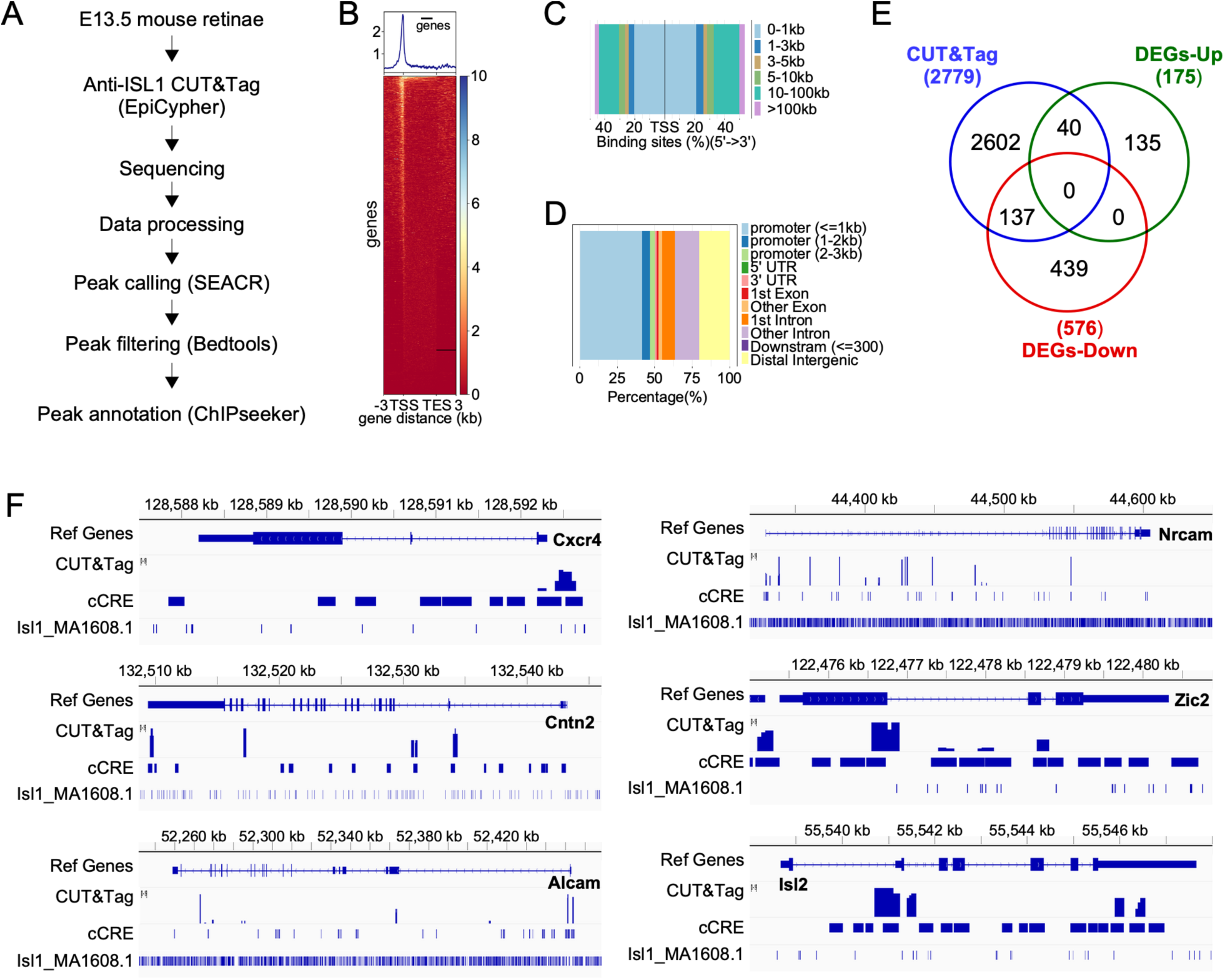
CUT&Tag analysis of ISL1-binding in the retina at E13.5. ***A,*** Workflow of CUT&Tag protocol. ***B,*** Genomic distribution of ISL1-bound peaks relative to the distance from transcriptional start site (TSS). TES indicates the transcriptional end site. ***C,*** Distribution of ISL1 binding loci relative to TSS. ***D,*** Genomic annotation of peaks identified by CUT&Tag analysis using antibodies against ISL1. ***E,*** Venn diagram showing the overlap of DEGs identified by RNA-Seq and genes found by CUT&Tag analysis. ***F,*** Integrative Genomics Viewer (IGV) track visualization of cis-regulatory elements (CRE) of selected genes bound by ISL1, identified by CUT&Tag analysis.

## Discussion

We have previously shown that loss of *Isl1* in the retina results in errors in axon pathfinding which are visible at the optic chiasm between E13.5 and E15.5: a delay in reaching the midline, presence of a smaller optic nerve, and axon fasciculation defects, and hypothesized that these axon growth defects could lead to the RGC cell death observed later, from E16.5 (Pan et al., 2008). In this study, by P0, we observe an aberrant, increased ipsilateral projection of RGC axons in the *Isl1-*null from regions of the retina that normally give rise only to contralateral RGCs. However, these mis-specified ipsilateral projections broadly target regions of the dorsal lateral geniculate nucleus and superior colliculus that are normally innervated by contralateral RGC axons. We show that ISL1 regulates the expression of several transcription factors, cell adhesion molecules and axon guidance factors that could control RGC laterality.

### Genetic programs controlling the development of contralateral RGCs

We show that the ZIC2^+^ RGC domain expands out of the ventrotemporal retina in the *Isl1-*null and *Zic2* expression levels are significantly upregulated in our *Isl1-*null RNA-seq data, suggesting that ISL1 could repress ZIC2 expression in developing RGCs. This is different from the *Isl2-*null retina, which displays increased expression of ZIC2 only within the VT region (Pak et al., 2004). The expansion of the ZIC2^+^ RGC domain in the *Isl1-*null is similar to that seen upon loss of *Pou3f1* in a recent study (Fries et al., 2023). *Pou3f1-*null mice also display increased ipsilateral projections at the optic chiasm and reduced contralateral RGCs. Some of the molecular players acting downstream of ISL1 and POU3F1 to regulate contralateral versus ipsilateral RGC growth appear to be similar, like the cell adhesion molecules *Alcam* and *Cntn2*. However, our RNA-Seq and gene expression analyses of control and *Isl1*-null retinae also reveal downregulation of genes implicated in the midline avoidance of ipsilateral RGC axons and the crossing of contralateral RGC axons such as *Shh*, *Nrcam*, and *Cxcr4* that appear unchanged in the transcriptome of the *Pou3f1-*null retina or upon POU3F1 overexpression in the retina (Fries et al., 2023). Furthermore, POU4F1, which has been reported to be essential for midline crossing of contralateral RGCs, and ISL2, a driver of the late born contralateral RGC projection that arises from the VT retina, are both direct transcriptional targets of ISL1 but not POU3F1 (this study and Pan et al., 2008; Fernández-Nogales et al., 2022; Fries et al., 2023). POU3F1 also upregulates expression of the SOXC family of transcription factors (SOX4, SOX11), which have been shown to regulate differentiation of contralateral RGCs by antagonizing Notch1-Hes5 signaling and facilitating contralateral RGC crossing at the midline by upregulating the expression of axon guidance receptors Plexin-A1 and Nr-CAM (Kuwajima et al., 2017). In comparison, our data show that ISL1 is required for the expression of Nr-CAM as well as the semaphorin receptor Neuropilin 1, but not Plexin-A1. Plexin-A1 and Nr-CAM are part of a receptor system on contralateral RGC axons required to overcome repulsive cues and cross the midline and the absence of either would result in a decrease in contralateral RGC crossing (Kuwajima et al., 2012). Contralateral RGC crossing is also enabled by Neuropilin 1 binding to its ligand VEGF-A which acts as an attractive cue at the optic chiasm (Erskine et al., 2011). Thus, ISL1 regulates RGC laterality through a set of unique downstream effectors as well as a few common ones, as other transcription factors. Interestingly, we found that *Nrcam* is also a direct transcriptional target of ISL1 in the inner ear in an earlier study which showed that loss of ISL1 results in defects in migration and axon pathfinding of sensory neurons of the inner ear (Xu et al., 2024).

### Eye specific targeting of brain nuclei is independent of laterality decisions at the midline

In the *Isl1-*null, RGCs arising from the non-VT retina mis-project ipsilaterally at the optic chiasm and nonetheless target regions of the dLGN and SC that are occupied by contralaterally projecting RGCs. These results demonstrate that target recognition is intrinsically encoded in RGCs in agreement with earlier studies of *EphB1* loss-of-function and *Zic2* gain-of-function. RGC axons from the VT retina of *EphB1-/-* mice are misrouted contralaterally at the OC but target the ipsilateral axon-recipient zone of the dLGN (Rebsam et al., 2009). Overexpression of ZIC2 in the center of the retina causes many RGCs to switch laterality and project ipsilaterally but these Zic2^+^ axons are found in regions of the dLGN and SC normally occupied by contralateral axons (García-Frigola and Herrera, 2010). However, refinement of these misrouted axons and achievement of finer eye specific segregation may still be disrupted as the molecular factors that control guidance at the midline may not be the same as those required for refinement within visual targets. For example, ZIC2 activates EPHB1, which controls axon laterality, and the serotonin transporter (SERT), which is dispensable for laterality decisions at the midline but required for axon refinement within visual targets (García-Frigola and Herrera, 2010).

## Acknowledgments

We thank Dr. Ling Pan for technical assistance. This study is supported by the NIH (R01DC021456 and R21EY035294) and the Georgia Research Alliance funding to L. Gan, and by Augusta University EM/Histology Core Module of the National Eye Institute Core grant (P30EY031631).

